# Collaborative hunting in artificial agents with deep reinforcement learning

**DOI:** 10.1101/2022.10.10.511517

**Authors:** Kazushi Tsutsui, Ryoya Tanaka, Kazuya Takeda, Keisuke Fujii

## Abstract

Collaborative hunting, in which predators play different and complementary roles to capture prey, has been traditionally believed as an advanced hunting strategy requiring large brains that involve high level cognition. However, recent findings that collaborative hunting have also been documented in smaller-brained vertebrates have placed this previous belief under strain. Here, we demonstrate that decisions underlying collaborative hunts do not necessarily rely on sophisticated cognitive processes using computational multi-agent simulation based on deep reinforcement learning. We found that apparently elaborate coordination can be achieved through a relatively simple decision process of mapping between observations and actions via distance-dependent internal representations formed by prior experience. Furthermore, we confirmed that this decision rule of predators is robust against unknown prey controlled by humans. Our results of computational ecology emphasize that collaborative hunting can emerge in various intra- and inter-specific interactions in nature, and provide insights into the evolution of sociality.

## Introduction

Cooperation among animals often provides fitness benefits to individuals in an competitive natural environment^1,2^. Cooperative hunting, in which two or more individuals engage in a hunt for the increase in successful prey capture, has been regarded as one of the most widely distributed forms of cooperation in animals^3^, and has received considerable attention because of the close links between cooperative behavior, its cognitive demand, and even sociality^4–7^. Cooperative hunts have been documented in a wide variety of species^7,8^, yet “collaboration” (or “collaborative hunting”), in which predators play different and complementary roles, has been reported in only a handful of vertebrate species^9–11^. For instance, previous studies have shown that mammals such as lions and chimpanzees are capable of dividing roles among individuals, such as chasing prey or blocking the prey’s escape path, to facilitate capture by the group^9,10^. The collaborative hunts appear to be achieved through elaborate coordination with other hunters, and is often believed as an advanced hunting strategy requiring large brains that involve high level cognition such as sharing intentions among predators or anticipating others’ movements^12^.

However, a growing number of recent findings have placed this previous belief under strain. That is, cases of intra- and inter-specific collaborative hunting have also been demonstrated in smaller-brained vertebrates such as birds^13^, reptiles^14^, and fish^15,16^, suggesting that collaborative hunting does not necessarily rely on complex cognitive processes. In other words, it seems possible that apparently elaborate hunting behavior can emerge in a relatively simple decision process in response to ecological needs^16^. However, the decision process underlying collaborative hunting remains poorly understood because most previous studies thus far have relied exclusively on field observations. Observational studies are essential for documenting such natural behavior, yet cannot, in principle, identify the specific decision process that results in coordinated behavior. This limitation arises because seemingly simple behavior can result from complex processes^17^ and vice versa^18^.

We therefore sought to further our understanding of the cognitive and decision processes underlying collaborative hunting by adopting a different approach, namely, computational multi-agent simulation based on deep reinforcement learning. Deep reinforcement learning mechanisms were originally inspired by animal associative learning^19^, and are thought to be closely related to neural mechanisms for reward-based learning centering on dopamine^20,21^. Given that associative learning is likely to be the most widely adopted learning mechanism in animals^22,23^, predator agents based on deep reinforcement learning may learn decision rules that result in collaborative hunting, if collaborative hunts do not necessarily require high level cognition.

Specifically, we first explored whether predator agents learn decision rules resulting in collaborative hunting and, if so, under what conditions through predator-prey interactions in a computational ecological environment. We then examined what internal representations mediate the decision rules. Furthermore, we confirmed the generality of the acquired predators’ decision rules using joint plays between agents (predators) and humans (prey). Our results showed that the acquisition of decision rules resulting in collaborative hunting is facilitated by a combination of two factors: prey capture difficulty of solitary hunting and food (i.e., reward) sharing following capture. We also found that the decisions underlying collaborative hunts were made through distance-dependent internal representations formed by prior experience. Furthermore, the decision rules robustly worked against unknown prey controlled by humans. These provide insights that collaborative hunts do not necessarily require high level cognition, and simple decision rules based on mappings between observations and actions can be practically useful in nature. Our approach of computational ecology bridges the gap between ecology, ethology, and neuroscience and may provides a comprehensive account of them.

## Results

We set out to model the decision process of predators and prey in an interactive environment. In this study, we focused on a chase and escape scenario in a two-dimensional open environment. Chase and escape may be complex phenomena in which two or more agents interact in environments, which change from moment to moment. Nevertheless, many studies have shown that the rules of chase/escape behavior (e.g., which direction to move at each time in a given situation) can be described by relatively simple mathematical models consisting of the current state (e.g., positions and velocities)^24–26^. We therefore considered modeling the agent’s decision process in a standard reinforcement learning framework for a finite Markov decision process, in which each sequence is a distinct state. In this framework, the agent interacts with the environment through a sequence of observations, actions, and rewards, and aims to select actions in a manner that maximizes cumulative future reward^27^.

We aimed to construct a biologically plausible (or considered to be more amenable to biological interpretation) simulation environment, and modeled an agent (predator/prey) with independent learning^28^ (Fig. 1a). Independent learning is one of the simplest forms of the multi-agent reinforcement learning algorithm, in which each agent treats the other agents as part of the environment and learns policies that are conditioned only on the agent’s local observation history. That is, in contrast to previous studies on multi-agent reinforcement learning^29–31^, our agents did not have access to models of the environment and observations and policies of other agents. For each agent *n*, the policy *π^n^* was represented by a neural network and optimized in the framework of deep Q-network^32^ (see Methods). The inputs to the neural network are subjectively available information about the environment and the outputs are the acceleration in 12 directions every 30 degrees in the relative coordinate system to the prey, which were determined based on previous findings on ecology^33^, ethology^24,25^, and neuroscience^34^. We assumed that delays in sensorimotor processing would be compensated for by estimation of the motion of self^35,36^ and others^37^, and the current information at each time step was taken as input as is. The predator(s) was rewarded for capturing the prey (+1), namely contacting the disks, and punished for moving out of the area (−1), and the prey was punished for being captured by the predator or for moving out of the area (−1). The time step was 0.1 s and the time limit in each episode was set to 30 s. The initial position of each episode was randomly selected from a range of −0.5 to 0.5 on the *x* and *y* axes.

**Figure 1.**
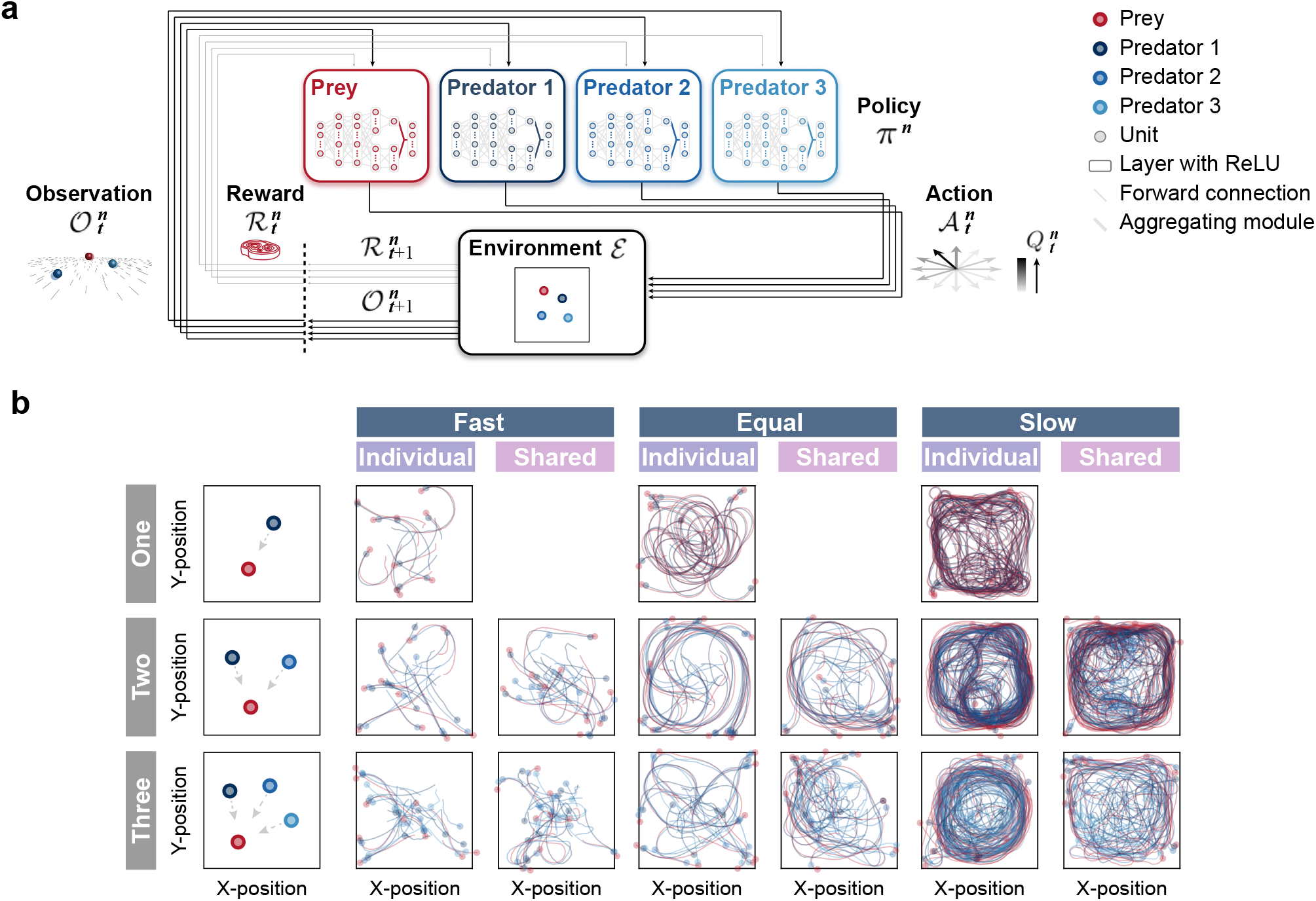
Agent architecture and examples of movement trajectories. (a) An agent’s policy is represented by a deep neural network (see Methods). An observation of the environment is given as input to the network. An action is sampled from the network’s output, and the agent receives a reward and the subsequent observation. The agent learns to select actions that maximizes cumulative future rewards. In this study, each agent learned its policy network independently, that is, each agent treats the other agents as part of the environment. This illustration shows a case with three predators. (b) The movement trajectories are examples of predator(s) (dark blue, blue, and light blue) and prey (red) interactions that overlay 10 episodes in each experimental condition. The experimental condition was set as the number of predators (one, two, and three), relative mobility (fast, equal, and slow), and reward sharing (individual and shared), based on ecological findings.

### Exploring the conditions under which collaborative hunting emerges

We first performed computational simulations with three experimental conditions to investigate under what conditions collaborative hunting emerges (Fig. 1b). As experimental conditions, we selected the number of predators, relative mobility, and prey (reward) sharing based on ecological findings^7,8^. For the number of predators, three conditions were set: 1 (one), 2 (two), and 3 (three). In all these conditions, the number of prey was set as 1. For the relative mobility, three conditions were set: 120% (fast), 100% (equal), and 80% (slow) for the acceleration exerted by the predator, based on that exerted by the prey. For the prey sharing, two conditions were set: with sharing (shared), in which all predators were rewarded when a predator catches the prey, and without sharing (individual), in which a predator was rewarded only when it catches prey by itself. In total, there were 15 conditions.

As the example trajectories show, under the fast and equal conditions, the predators often caught their prey shortly after starting the episode, whereas under the slow condition, the predators relatively struggled to catch their prey (Fig. 1b). To evaluate their behavior, we calculated the proportion of successful predation and mean episode duration. For the fast and equal conditions, predations were successful in almost all episodes, regardless of the number of predators and the presence or absence of reward sharing (e.g., 0.99 ± 0.00 for the one × fast and one × equal conditions; Fig. S1 top). This indicates that in situations where predators were faster than or equal in speed to their prey, they almost always succeeded in capturing the prey, even when they were alone. Although the mean episode duration decreased with an increasing number of predators in both fast and equal conditions, the difference was small (Fig. S1 bottom). As a whole, these results indicate that there is little benefit of cooperating among multiple predators in the fast and equal conditions. As it is unlikely that cooperation among predators will emerge under such conditions in nature from an evolutionary perspective^1,2^, the analysis below is limited to the slow condition. For the slow condition, a solitary predator was rarely successful, and the proportion of successful predation increased with the number of predators (Fig. 2a top). Moreover, the mean duration decreased with increasing number of predators (Fig. 2a bottom). These results indicate that, under the slow condition, the benefits of cooperation among multiple predators are significant. In addition, except for the two × individual condition, the increase in the proportion of success with increasing number of predators was much greater than the theoretical prediction^3^, calculated based on the proportion of solitary hunting, assuming that each predator’s performance is independent of the others’ (see Methods). These results indicate that under these conditions, elaborate hunting behavior (e.g., “collaboration”) that is qualitatively different from hunting alone may emerge.

**Figure 2.**
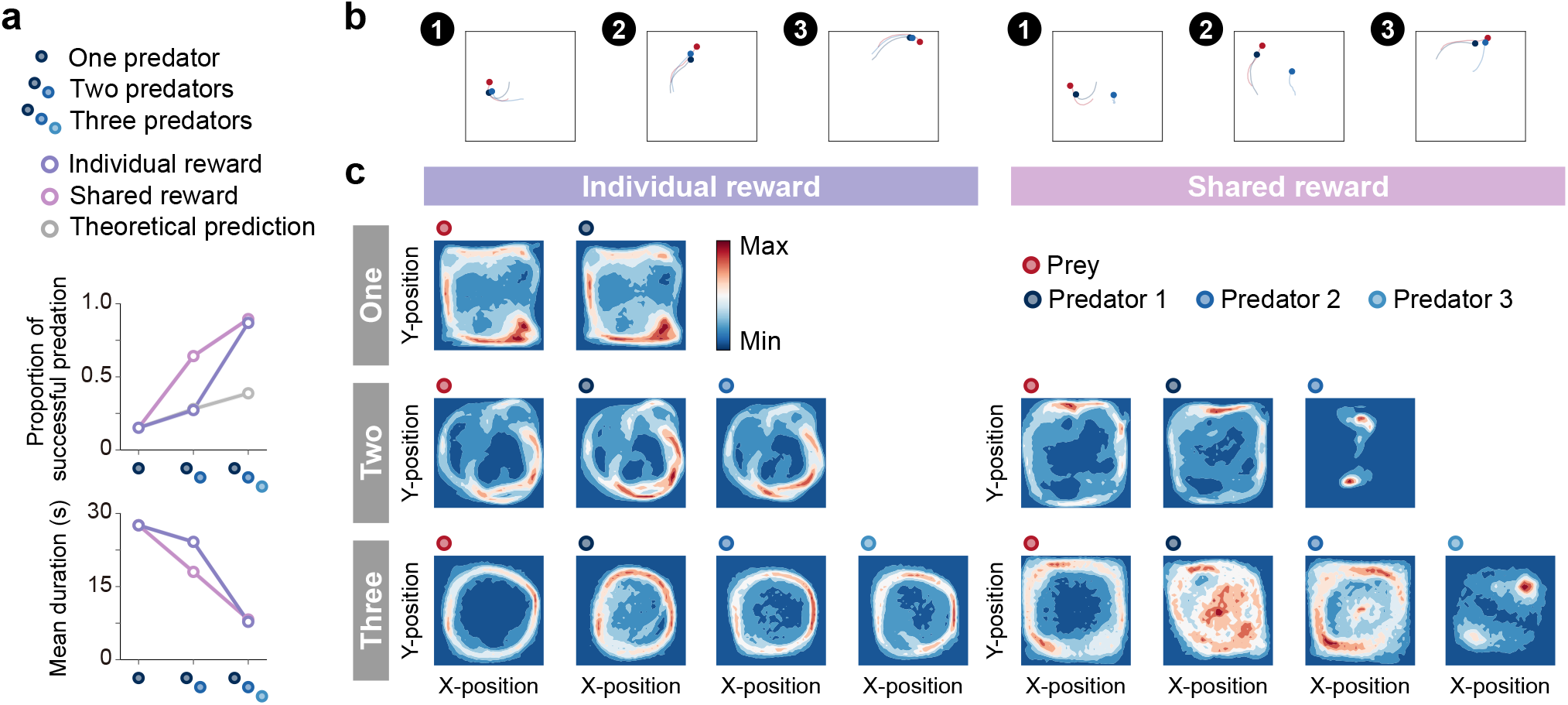
Emergence of collaborations among predators. (a) Proportion of successful predation (top) and mean episode duration (bottom). For both panels, quantitative data denote the mean of 100 episodes ± s.e.m across 10 random seeds. The error bars are barely visible because the variation is negligible. The theoretical prediction values were calculated based on the proportion of solitary hunts (see Methods). The proportion of successful predation increased as the number of predators increased (*F*_number(2,18)_ = 1346.67, *p* < 0.001; *η*^2^ = 0.87; one vs. two: *t*_(9)_ = 20.38, *p* < 0.001; two vs. three: *t*_(9)_ = 38.27, *p* < 0.001). The mean duration decreased with increasing number of predators (*F*_number(2,18)_ = 1564.01, *p* < 0.001; *η*^2^ = 0.94; one vs. two: *t*_(9)_ = 15.98, *p* < 0.001; two vs. three: *t*_(9)_ = 40.65, *p* < 0.001). (b) Typical example of different predator routes between the individual (left) and shared (right) conditions, in the two-predator condition. The numbers (1 to 3) show a series of state transitions (every second) starting from the same initial position. Each panel shows the agent positions and the trajectories leading up to that state. In these instances, the predators eventually failed to capture the prey within the time limit (30 s) under the individual condition, whereas the predators successfully captured the prey in only 3 s under the shared condition. (c) Comparison of heat maps between individual (left) and shared (right) reward conditions. The heat map of each agent was made based on the frequency of stay in each position, which is cumulative for 1,000 episodes (100 episodes × 10 random seeds). In the individual condition, there were relatively high correlations between the heat maps of the prey and each predator, regardless of the number of predators (One: *r* = 0.95, *p* < 0.001, Two: *r* = 0.83, *p* < 0.001 in predator 1, *r* = 0.78, *p* < 0.001 in predator 2, Three: *r* = 0.41, *p* < 0.001 in predator 1, *r* = 0.56, *p* < 0.001 in predator 2, *r* = 0.45, *p* < 0.001 in predator 3). In contrast, in the shared condition, only one predator had a relatively high correlation, whereas the others had low correlations (Two: *r* = 0.65, *p* < 0.001 in predator 1, *r* = 0.01, *p* = 0.80 in predator 2, Three: *r* = 0.17, *p* < 0.001 in predator 1, *r* = 0.54, *p* < 0.001 in predator 2, *r* = 0.03, *p* = 0.23 in predator 3).

Then, we examined agent behavioral patterns and found that there were differences in the movement path that predators take to catch their prey among the conditions (Fig. 2b). As shown in the typical example, under the individual condition, both predators moved similarly toward their prey (Fig. 2b left) and, in contrast, under the shared condition, one predator moved toward their prey while the other predator moved along a different route (Fig. 2b right). To ascertain their behavioral patterns, we created heat maps showing the frequency of agent presence at each location in the play area (Fig. 2c). We found that there was a noticeable difference between the individual and shared reward conditions. That is, in the individual condition, the heat maps of prey and respective predators were quite similar (Fig. 2c left), whereas this was not always the case in the shared condition (Fig. 2c right). In particular, the heat maps of predator 2 in the two-predator condition and predator 3 in the three-predator condition showed localized concentrations (Fig. 2c far right, respectively). To assess these differences among predators in more detail, we compared the predators’ decisions (i.e., action selections) in these condition with that in the one-predator condition (i.e., solitary hunts) using two indices, concordance rate and circular correlation^38^ (Fig. S2). Following previous studies^39^, we also calculated the ratios of distance moved during hunting among predators (Fig. S3). As a whole, these buttress that predators with similar heat maps to the prey behaved as ‘chaser’ (or ‘driver’), while predators with different heat maps to the prey behaved as ‘blocker’ (or ‘ambusher’). That is, our results show that, although most predators acted as chasers, some predators were acted as blockers rather than chasers in the shared condition, indicating the emergence of collaborative hunting characterized by role divisions among predators under the condition.

### Searching internal representation underlying collaboration

We next sought the predators’ internal representations to better understand how such collaborative hunting is accomplished. Using a two-dimensional t-distributed stochastic neighbor embedding (t-SNE)^40^, we visualized the last hidden layers of the state and action streams in the policy network as internal representations of agents (Fig. 3 and Figs. S5 to S7). To understand how each agent represents its environment and what aspects of the state are well represented, we examined the relationship between the scenes of a typical scenario and their corresponding points on the embedding (Figs. 3a and b). As expected, when the predator is likely to catch its prey (e.g., the scene 4), the predator estimated a higher state value, whereas, when the predator is not (e.g., the scene 5), the predator estimated a lower state value (Fig. 3a top). Related to this, the variance of action values tends to be larger for both predator and prey when they are close (Fig. 3a bottom), indicating that the difference in the value of choosing each action is greater when the choice of action is directly related to the reward (see also Fig. S4). These results suggest that the agents were able to learn the networks that outputs the estimations of state and action values consistent with our intuition.

**Figure 3.**
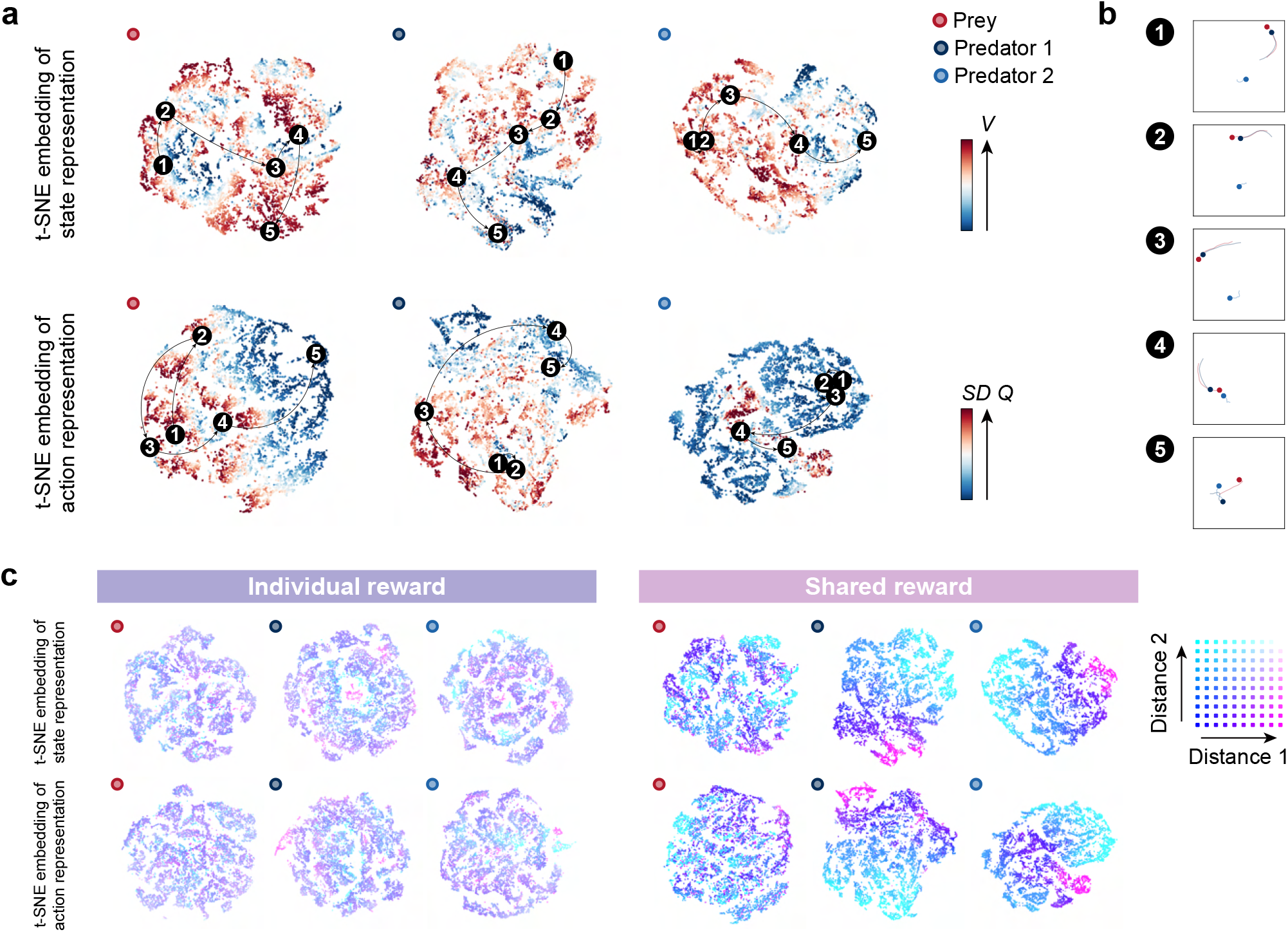
Embedding of internal representations underlying collaborative hunting. (a) Two-dimensional t-SNE embedding of the representations in the last hidden layers of the state-value stream (top) and action-value stream (bottom) in the shared reward condition. The representation is assigned by the policy network of each agent to states experienced during predator-prey interactions. The points are colored according to the state values and standard deviation of the action values, respectively, predicted by the policy network (ranging from dark red (high) to dark blue (low)). (b) Corresponding states for each number in each embedding. The number (1 to 5) in each embedding corresponds to a selected series of state transitions. The series of agent positions in the state transitions (every second) and, for ease of visibility, the trajectories leading up to that state are shown. (c) Embedding colored according to the distances between predators and prey in the individual (left) and shared (right) reward conditions. Distance 1 and 2 denote the distance between predator 1 and prey and predator 2 and prey, respectively. If two distances are short, the point is colored blue; if long, it is colored white).

Furthermore, we found a distinct feature in the embedding of predators’ representations. Specifically, in certain state transitions, the position of the points on the embedding changed little, even though the agents were moving (e.g., the scenes 1 to 2 on the embedding of the predator 2). From this, we reasoned that predators’ representations may be encoded how far away the others are from them than about where themselves and the others are. To test our reasoning, we colored the representations according to the distance between predators and prey; distance 1 denotes the distance between predator 1 and prey, and distance 2 denotes that between predator 2 and prey. As a result, the representations of predators in the shared condition could be clearly separated by the distance-dependent coloration (Fig. 3c right), in contrast to those in the individual condition (Fig. 3c left). These indicate that the predators in the shared condition estimated state and action values and made decisions, mediated by distance-dependent representations (see Fig. S6 for prey’s decision).

### Evaluating the playing strength of predator agents using joint play with humans

Finally, to verify the generality of predators’ decisions against unknown prey, we conducted an experiment of joint play between agents and humans. In the joint play, human participants controlled an prey on a screen with the joystick of a controller. The objective, as in the above computational simulation, was to escape until the end of the episode (30 s) without being caught by the predators and without leaving the area. We found that the outcomes of the joint play show similar trends to those of the computer simulation (Fig. 4a). That is, the proportion of successful predation increased and the mean episode duration decreased, as the number of predators increased. These indicate that the predator agents’ decision rules worked well for the prey controlled by humans. To visualize the associations of states experienced by predator agents versus agents and versus humans, we show colored two-dimensional t-SNE embedding of the representations in the last hidden layer of the state stream (Fig. 4b and Fig. S8). These showed that, in contrast to a previous study^32^, the states were quite separate, suggesting that predator agents experienced unfamiliar states when playing against the prey controlled by humans. This unfamiliarity may make it difficult for predators to make proper decisions. Indeed, in the one-predator condition, the predator agent occasionally exhibited suspicious behavior (e.g., staying in one place; see Fig. S9). On the other hand, in the two- and three-predator conditions, predator agents rarely exhibited such behavior and showed superior performance. This indicates that decision rules of cooperative hunting acquired in certain environments could be applied in other somewhat different environments.

**Figure 4.**
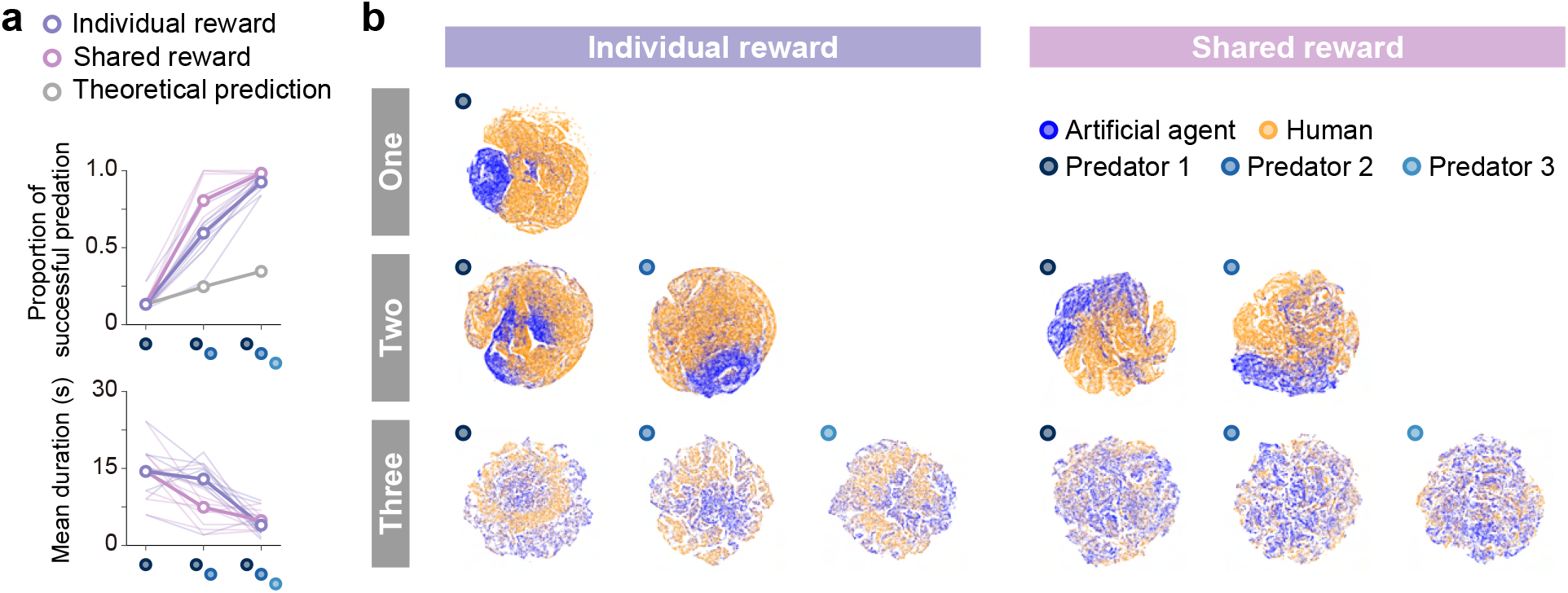
Superior performance of predator agents for prey controlled by humans and comparison of internal representations. (a) Proportion of successful predation (top) and mean episode duration (bottom). For both panels, the thin line denotes the performance for each participant, and the thick line denotes the mean. The theoretical prediction values were calculated based on the mean of proportion of solitary hunts. The proportion of successful predation increased as the number of predators increased (*F*_number(1.28,11.48)_ = 276.20, *p* < 0.001; *η*^2^ = 0.90; one vs. two: *t*_(9)_ = 13.80, *p* < 0.001; two vs. three: *t*_(9)_ = 5.9402, *p* < 0.001). The mean duration decreased with increasing number of predators (*F*_number(2,18)_ = 23.77, *p* < 0.001; *η*^2^ = 0.49; one vs. two: *t*_(9)_ = 2.60, *p* = 0.029; two vs. three: *t*_(9)_ = 5.44, *p* < 0.001). (2) Comparison of two-dimensional t-SNE embedding of the representations in the last hidden layers of state-value stream between self-play (predator agents vs. prey agent) and joint play (predator agents vs. prey human).

## Discussion

It has long been thought that collaborative hunting is an advanced hunting strategy requiring large brains that involve high level cognition. This traditional view have held that anticipations of the other hunters and prey would be essential for the collaboration. Here we have shown that the “collaboration”^10^ can emerge in group hunts of artificial agents in the absence of explicit predictive or planning mechanisms. This means that, in contrast to the traditional view, predators do not need to be able to anticipate the future movements of the other predators and prey for the collaboration and that apparently elaborate coordination can be accomplished by a relatively simple decision rules, that is, mappings between observations and actions. This result advances our understanding of cooperative hunting behavior and its decision process, and may offer a novel perspective on the evolution of sociality.

Our results on agent behavior are broadly consistent with previous studies on observations of animal behavior in nature. First, as the number of predators increased, success rates increased and hunting duration decreased^5^. Second, whether collaborative hunts emerge depended on two factors: the success rate of hunting alone^12,41^ and the presence or absence of meat (food) sharing following prey capture^42,43^. Third, in repetition of collaborative hunts, the role of each predator was basically fixed, but could be flexibly interchanged^9,12^. Finally, predator agents in this study acquired different strategies depending on the conditions despite having exactly the same initial values (i.e., network weights), resonating with the findings that lions and chimpanzees living in different regions exhibit different hunting strategies^9,44^. These results suggest the validity of our computational simulations and highlight the close link between predators’ behavioral strategies and their living environments.

We found that the mappings resulting in collaborative hunting were mediated by distance-dependent internal representations. Deep reinforcement learning have held promise for providing a comprehensive framework for studying the interplay among learning, representation, and decision making^45,46^, but such efforts for natural behavior have been limited^47,48^. Our result that distance-dependent representations mediate collaborative hunting is reminiscent of a recent idea about the decision rules obtained by observation in fish^16^, suggesting that notable parallels on decision making between artificial agents and animals. Notably, a predator’s subjective input variables do not include variables corresponding to the distance(s) between the other predator(s) and prey. This means that the predators in the shared conditions acquired the internal representation relating with distance to prey, which would be a geometrically reasonable indicator, by optimization through interaction with their environment.

The predator agents’ decision rules (i.e., policy networks) acquired through interactions with other agents (i.e., self-play) were also useful for human-controlled, unknown, prey, despite the dissociation of the experienced states. This suggests that decisions rules formed by associative learning can successfully address natural problems, such as catching prey with somewhat different movement patterns than one’s usual prey. Note that the learning mechanism of associative learning (or reinforcement learning) is relatively simple, but it allows for flexible behavior in response to situations, in contrast to innate and simple stimulus-response. Indeed, our agents generated complex trajectories, and the prey agents outperformed the prey controlled by humans in the proportion of successful escape. Our view that decisions for successful hunting are made through representations formed by prior experience is a counterpart to the recent idea that computational relevance for successful escape may be cached and ready to use, instead of being computed from scratch on the spot^17^. Suppose that the decision process of animals in predator-prey dynamics adopts such a manner, it may be a product of natural selection that allows for rapid, robust, and flexible action generation in strongly time-constrained interactions.

In conclusion, we demonstrated that the decisions underlying collaborative hunting among artificial agents can be achieved through a mapping between observation and action mediated by representations formed by prior experience. This means that collaborative hunting can emerge in the absence of highly cognitive mechanisms such as anticipating others’ movements or sharing the intentions among predators, supporting the recent idea that collaborative hunting does not necessarily rely on complex cognitive processes in brains. Our computational ecology is an idealization of a real predator-prey environment. Given that chase and escape often involve various factors, such as energy cost^49^, signal communication^50^, and local surroundings^17^, these results are only a first step on the path to understanding real decisions in predator-prey dynamics. Because of the notable parallel between artificial agents’ and animals’ behavior, the computational ecological environment seems particularly well positioned to understand real-world decisions for survival. Although animals’ bodies and nervous systems should have evolved in close association to increase their chances of survival in a given environment, few studies have linked behavior, decision making, and neural computation. Thus, we believe that our results provide a useful advance toward understanding natural value–based decisions and forge a critical link between ecology, ethology, and neuroscience.

## Methods

### Environment

The predator and prey interacted in a two-dimensional world with continuous space and discrete time. This environment was constructed by modifying an environment called the predator-prey in multi-agent particle environment^31^. Specifically, the position of each agent was calculated by integrating the acceleration (i.e., selected action) twice with the Euler method, and viscous resistance proportional to velocity was considered. The modifications were that the action space (play area size) was constrained to the range of −1 to 1 on the *x* and *y* axes, all agent (predator/prey) disk diameters were set to 0.1, landmarks (obstacles) were eliminated, and predator-to-predator contact was ignored for simplicity. The predator(s) was rewarded for capturing the prey (+1), namely contacting the disks, and punished for moving out of the area (−1), and the prey was punished for being captured by the predator or for moving out of the area (−1). The predator and prey were represented as a red and blue disk, respectively, and the play area was represented as a black square surrounding them. The time step was 0.1 s and the time limit in each episode was set to 30 s. The initial position of each episode was randomly selected from a range of −0.5 to 0.5 on the *x* and *y* axes. If the predator captured the prey within the time limit, the predator was successful, otherwise, the prey was successful. If one side (predators/prey) moved out of the area, the other side (prey/predators) was deemed successful.

### Experimental condition

We selected the number of predators, relative mobility, and prey (reward) sharing as experimental conditions, based on ecological findings^7,8^. For the number of predators, three conditions were set: 1 (one), 2 (two), and 3 (three). In all these conditions, the number of preys was set as 1. For the relative mobility, three conditions were set: 120% (fast), 100% (equal), and 80% (slow) for the acceleration exerted by the predator, based on that exerted by the prey. For the prey sharing, two conditions were set: with sharing (shared), in which all predators were rewarded when a predator catches the prey, and without sharing (individual), in which a predator was rewarded only when it catches prey by itself. In total, there were 15 conditions.

### Agent architecture

We considered a sequential decision-making setting in which a single agent interacts with an environment 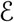 in a sequence of observations, actions, and rewards. At each time-step *t*, the agent observes a state 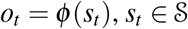 and selects an action *a_t_* from a discrete set of actions 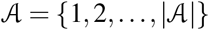. One time step later, in part as a consequence of its action, the agent receives a reward, 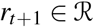, and moves itself to a new state *s*_*t*+1_. In the MDP, the agent thereby gives rise to a sequence that begins like this: *s*_0_, *a*_0_, *r*_1_, *s*_1_, *a*_1_, *r*_2_, *s*_2_, *a*_2_, *r*_3_,…, and learn behavioral rule (policy) that depend upon these sequences.

The goal of the agent is to maximize the expected discounted return over time through its choice of actions^27^. The discounted return *R_t_* was defined as 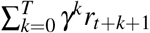, where *γ* ∈ [0, 1] is a parameter called the discount rate that determines the present value of future rewards, and *T* is the time step at which the task terminates. The state-value function, action-value function, and advantage function are defined as 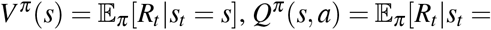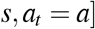, and *A^π^* (*s*, *a*) = *Q^π^* (*s*, *a*) − *V^π^* (*s*), respectively, where *π* is a policy mapping states to actions. The optimal action-value function *Q*^⋆^(*s*, *a*) is then defined as the maximum expected discounted return achievable by following any strategy, after observing some state *s* and then taking some action *a*, 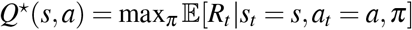. The optimal action-value function can be computed recursively obeying the Bellman equation:

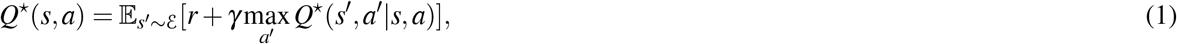

where *s*′ and *a*′ are the state and action at the next time-step, respectively. This is based on the following intuition: if the optimal value *Q*^⋆^(*s*′, *a*′) of the state *s*′ was known for all possible actions *a*′, the optimal strategy is to select the action *a*′ maximizing the expected value of *r* + *γ* max_*a*′_ *Q*^⋆^(*s*′, *a*′). The basic idea behind many reinforcement learning algorithms is to estimate the action-value function by using the Bellman equation as an iterative update; 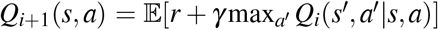. Such value iteration algorithms converge to the optimal action-value function in situations where all states can be sufficiently sampled, *Q_i_* → *Q*^⋆^ as *i* → ∞. In practice, however, it is often difficult to apply this basic approach, which estimates the action-value function separately for each state, to real-world problems. Instead, it is common to use a function approximator to estimate the action-value function, *Q*(*s*, *a*; *θ*) ≈ *Q*^⋆^(*s*, *a*).

There are several possible ways of function approximation, yet we here use a neural network function approximator referred to as deep *Q*-network (DQN)^32^ and some of its extensions to overcome the limitations of the DQN, namely double DQN^51^, prioritized experience replay^52^, and dueling network^53^. Deep *Q*-network is the first deep learning model to successfully learn agents’ policies directly from high-dimensional sensory input using reinforcement learning^32^. Naively, a *Q*-network with weights *θ* can be trained by minimizing a loss function 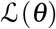 that changes at each iteration *i*,

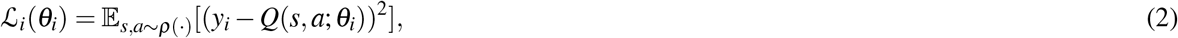

where *y_i_* = *r* + *γ* max_*a*′_ *Q*(*s*′, *a*′; *θ*_*i*−1_|*s*, *a*) is the target value for iteration *i*, and *ρ*(*s*, *a*) is a probability distribution over states *s* and actions *a*. The parameters from the previous iteration *θ*_*i*−1_ are kept constant when optimizing the loss function 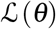. By differentiating the loss function with respect to the weights we arrive at the following gradient,

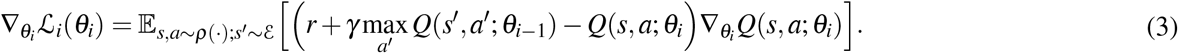

We could attempt to use the simplest *Q*-learning to learn the weights of the network *Q*(*s*, *a*; *θ*) online; however, this estimator performs poorly in practice. In this simplest form, they discard incoming data immediately, after a single update. This results in two issues: (i) strongly correlated updates that break the i.i.d. assumption of many popular stochastic gradient-based algorithms and (ii) the rapid forgetting of possibly rare experiences that would be useful later. To address both of these issues, a technique called experience replay is often adopted^54^, in which the agent’s experiences at each time-step *e_t_* = (*s_t_*, *a_t_*, *r*_*t*+1_, *s*_*t*+1_) are stored into a data-set (also referred to as replay memory) 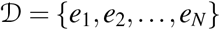, where *N* is the data-set size, for some time. When training the *Q*-network, instead of only using the current experience as prescribed by standard *Q*-learning, mini-batches of experiences are sampled from 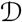 uniformly at random to train the network. This enables breaking the temporal correlations by mixing more and less recent experiences for the updates, and rare experiences will be used for more than just a single update. Another technique, called as target-network, is also often used for updating to stabilize learning. To achieve this, the target value *y_i_* is replaced by 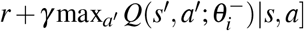, where 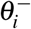 are the weights frozen for a fixed number of iterations. The full algorithm combining these ingredients, namely experience replay and target-network, is often called a deep Q-network (DQN), and its loss function takes the form:

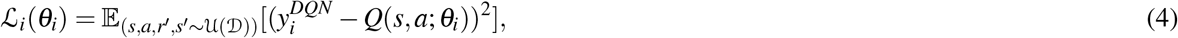

where

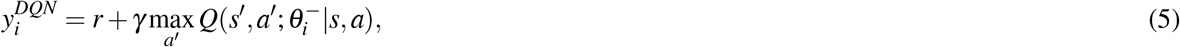

and 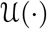 is a uniform sampling.

It has been known that a *Q*-learning algorithm performs poorly in some stochastic environments. This poor performance is caused by large overestimations of action values. These overestimations result from a positive bias that is introduced because *Q*-learning uses the maximum action value as an approximation for the maximum expected action value. As a method to alleviate the performance degradation due to the overestimation, double *Q*-learning, which applies the double estimator, was proposed^55^. Double DQN (DDQN) is an algorithm that applies the double *Q*-learning method to DQN^51^. For the DDQN, the maximum operation in the target is decomposed into action selection and action evaluation, and the target value in the loss function (i.e., Eq. 5) for iteration *i* is replaced as follows:

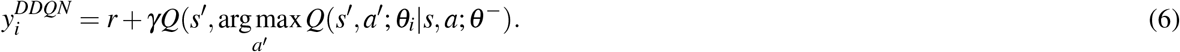

Prioritized experience replay is a method that aims to make the learning more efficient and effective than if all transitions were replayed uniformly^52^. For the prioritized replay, the probability of sampling from the data-set for transition *i* is defined as

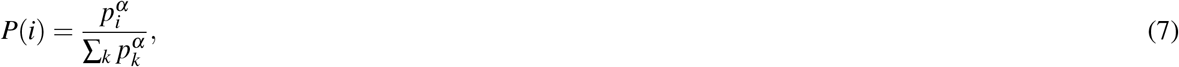

where *p_i_* > 0 is the priority of transition for iteration *i* and the exponent *α* determines how much prioritization is used, with *α* = 0 corresponding to uniform sampling. The priority *p_i_* is determined by *p_i_* = |*δ_i_*| + *ε*, where *δ_i_* is a temporal-difference (TD) error (e.g., 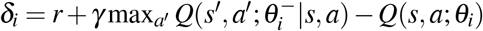 in DQN) and *ε* is a small positive constant that prevents the case of transitions not being revisited once their error is zero. Prioritized replay introduces sampling bias, and therefore changes the solution to which the estimates will converge. This bias can be corrected by importance-sampling (IS) weights 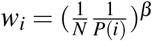 that fully compensate for the non-uniform probabilities *P*(*i*) if *β* = 1.

Dueling network is a neural network architecture designed for value-based algorithms such as DQN^53^. This features two streams of computation, the value and advantage streams, sharing a common encoder, and is merged by an aggregation module that produces an estimate of the state-action value function. Intuitively, we can expect the dueling network to learn which states are (or are not) valuable, without having to learn the effect of each action for each state. For the reason of stability of the optimization, the last module of the network is implemented as follows:

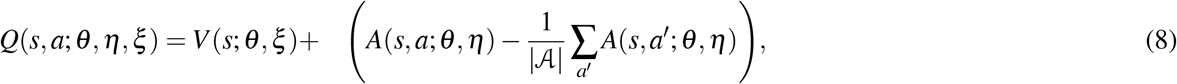

where *θ* denotes the parameters of the common layers, whereas *η* and *ξ* are the parameters of the layers of the two streams, respectively. Although this loses the original semantics of *V* and *A*, subtracting the mean helps with identifiability while preserving the stability of the optimization. In addition, it does not change the relative rank of the *A* (and hence *Q*) values, and suffices to evaluate the advantage stream to make decisions. It has been experimentally shown that this module works well in practice.

We here aimed to construct a biologically plausible (or considered to be more amenable to biological interpretation) simulation environment, and modeled an agent (predator/prey) with independent learning (IL)^28^. IL is one of the simplest forms of MARL, in which each agent treats the other agents as part of the environment and learns policies that are conditioned only on an agent’s local observation history. That is, in contrast to previous studies on multi-agent reinforcement learning^29–31^, our agents did not have access to models of the environment and observations and policies of other agents. For each agent *n*, the policy *π^n^* is represented by a neural network and optimized, with the framework of DQN including DDQN, prioritized replay, and dueling architecture. The loss function of each agent takes the form:

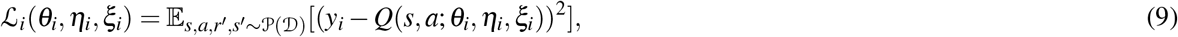

where

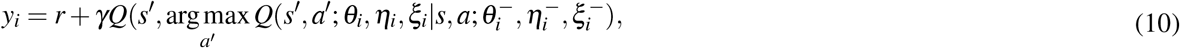

and 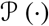 is a prioritized sampling. For simplicity, we omitted the agent index *n* in these equations.

### Training details

The neural network is composed of four layers. There is a separate output unit for each possible action, and only the state representation is an input to the neural network. The inputs to the neural network are the positions of oneself and others in the absolute coordinate system (*x*- and *y*-positions) and the positions and velocities of oneself and others in the relative coordinate system (*u*- and *v*-positions and *u*- and *v*-velocities) (see Fig. S10), which are determined based on findings in ethology^24,25^ and neuroscience^34^. We assumed that delays in sensory processing are compensated for by estimation of motion of self^35,36^ and others^37^ and the current information at each time was used as input as is. The outputs are the acceleration in 12 directions every 30° in the relative coordinate system, which are determined with reference to an ecological study^33^. After the first two hidden layers of the MLP with 64 units, the network branches off into two streams. Each branch has one MLP layer with 32 hidden units. ReLU was used as the activation function for each layer^56^. The network parameters *θ^n^*, *η^n^*, and *ξ*^*n*^ were iteratively optimized via stochastic gradient descent with the Adam optimizer^57^. In the computation of the loss, we used Huber loss to prevent extreme gradient updates^58^. The model was trained for 10^6^ episodes, and the network parameters were copied to the target-network every 2000 episodes. The replay memory size was 10^4^, the minibatch size during training was 32, and the learning rate was 10^−6^. The discount factor *γ* was set to 0.9, and *α* was set to 0.6. We used an *ε*-greedy policy as the behavior policy *π^n^*, which chooses a random action with probability *ε* or an action according to the optimal *Q* function 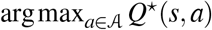 with probability 1 − *ε*. In this study, *ε* was annealed linearly from 1 to 0.1 over the first 10^4^ episodes and fixed at 0.1 thereafter.

### Evaluation

The model performance was evaluated using the trained model. The initial position and termination criteria in each episode were the same as in training. During the evaluation, *ε* was set to 0, and each agent took greedy actions. We first conducted a computational experiment (self-play: predator agent vs. prey agent) and then conducted a human behavioral experiment (joint play: predator agent vs. prey human). In the computational experiment, we simulated 100 episodes for each of the 10 random seeds(i.e., different initial positions), for a total of 1000 episodes in each condition. In the joint play, human participants controlled prey on a screen using the joystick of a controller and interacted with the predator agents for 50 episodes in each condition.

### Participants

Ten males participated in the experiment (aged 22-25, mean = 23.5, s.d. = 1.2). All participants were right-handed but one, had normal or corrected-to-normal vision, and were naïve to the purpose of the study. This study was approved by the Ethics Committee of Nagoya University Graduate school of Informatics. Informed consent was obtained from each participant before the experiment. Participants received 1,000 yen per hour as a reward.

### Apparatus

Participants were seated in a chair, and they operated the joystick of an Xbox One controller that could tilt freely in any direction to control a disk on the screen. The stimuli were presented on a 26.5-inch monitor (EIZO EV2730Q) at a refresh rate of 60 Hz. A gray square surrounding the disks was defined as the play area. The diameter of each disk on the screen was 2.0 cm, and the width and height of the area were 40.0 cm. The acceleration of each disk on the screen was determined by the inclination of the joystick on the controller. Specifically, the acceleration was added when the degree of joystick tilt exceeded half of the maximum tilt, and the direction of the acceleration was selected from 12 directions discretized every 30 degrees in an absolute coordinate system corresponding to the direction of joystick tilt. The reason for setting the direction of acceleration with respect to the absolute coordinate system, rather than the relative coordinate system, in the human behavioral experiment was to allow participants to control more intuitively. The position and velocity of each disk on the screen were updated at 10 Hz (corresponding to the computational simulation) and the position during the episodes was recorded at 10 Hz on a computer (MacBook Pro) with Psychopy version 3.0. The viewing distance of the participants was about 60 cm.

### Design

Participants controlled a red disk representing the prey on the screen. They were asked to escape from the predator for 30 seconds without leaving the play area. The agent’s initial position and the outcome of the episode were determined as described above. The experimental block consisted of 5 sets of 10 episodes, with a warm-up of 10 episodes to get used to the task. In this experiment, we focused on the slow condition and there were thus five experimental conditions (one, two × individual, two × shared, three × individual, and three × shared). Each participant played one block (i.e., 50 episodes) of each experimental condition. The order of the experimental conditions was pseudo-randomized across participants.

### Data analysis

All data analysis was performed in Python 3.7. Successful predation was defined as the sum of the number of predators catching prey and the number of prey leaving the play area. The theoretical prediction assumes that each predator’s performance is independent of the others’ performance, and was defined as follows:

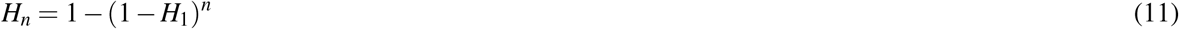

where *H_n_* and *H*_1_ denote the proportion of successful predation when the number of predators is *n* and 1, respectively. The duration was defined as the time from the beginning to the end of the episode, with a maximum duration of 30 s. The heat maps was made based on the frequency of stay in each position, which was divided the play area into 1600 (40 × 40). The concordance rate was calculated by comparing the actual selected action by each agent in the two or three conditions with the action that would be chosen by the agent in the one condition if it were placed in a same situation. The circular correlation coefficient was calculated by converting the selected actions (1 to 12) into angles (0 to 330 degrees)^38^, and in this analysis, action 13 (do nothing) was excluded from the analysis. The two-dimensional embedding was made by transforming the vectors in the last hidden layers of state-value stream and action-value stream in the policy network using t-distributed stochastic neighbor embedding (t-SNE)^40^ (see Fig. S11). To reduce the influence of extremely large or small values, the color ranges of *V* value, SD *Q* value, and distance were limited from the 5th percentile to the 95th percentile of whole values experienced by each agent (see Figs. S12 to S14).

### Statistics

All quantitative data are reported as mean ± s.e.m. across random seeds in the computational experiment and across participants in the human experiment. In the human experiment, sample sizes were not predetermined statistically, but rather were chosen based on field standards. The data were analyzed using one- or two-way repeated-measures analysis of variance (ANOVA) as appropriate. For these tests, Mauchly’s test was used to test sphericity; if the sphericity assumption was violated, degrees of freedom were adjusted by the Greenhouse–Geisser correction. *p* values were adjusted by the Holm–Bonferroni method for multiple comparisons. The data distribution was assumed to be normal for multiple comparisons, but this was not formally tested. Two-tailed statistical tests were used for all applicable analyses. The significance level was set at an alpha value of 0.05. The theoretical prediction was excluded from statistical analyses (Figs. 2a and 4a) because it is obvious that the proportion of successful predation increases as the number of predators increases from the equation. Specific test statistics, *p* values, and effect sizes for the analyses are detailed in the corresponding figure captions. All statistical analyses were performed using R version 4.0.2 (The R Foundation for Statistical Computing).

## Data availability

The datasets generated and analyzed in this study are available from the corresponding authors on reasonable request.

## Code availability

The codes used for computational simulation and data analysis are available from the corresponding authors on reasonable request.

## Acknowledgements

This work was supported by JSPS KAKENHI (Grant Numbers 20H04075, 21H04892, 21H05300, and 22K17673) and JST PRESTO (JPMJPR20CA).

## Author contributions

K.Ts. and K.F. conceived the study. K.Ts. developed and implemented the software. K.Ts. and K.F. designed the experiments. K.Ts. conducted the experiments. K.Ts. analyzed the data. K.Ts., R.T., K.Ta, and K.F. wrote the manuscript.

## Additional information

Correspondence and requests for materials should be addressed to Kazushi Tsutsui.

