## Supplementary Figures for "Collaborative hunting in artificial agents with deep reinforcement learning"

**This PDF file includes:**

Fig. S1 to S14

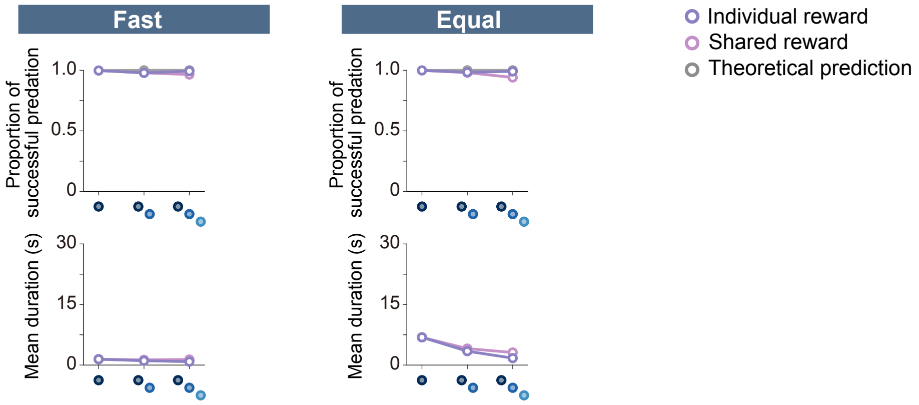


**Figure S1.** Proportion of successful predation (top) and mean episode duration (bottom) in the fast and equal conditions. For both panels, quantitative data denote the mean of 100 episodes ± s.e.m across 10 random seeds. The theoretical prediction values were calculated based on the proportion of solitary hunts (see Methods).

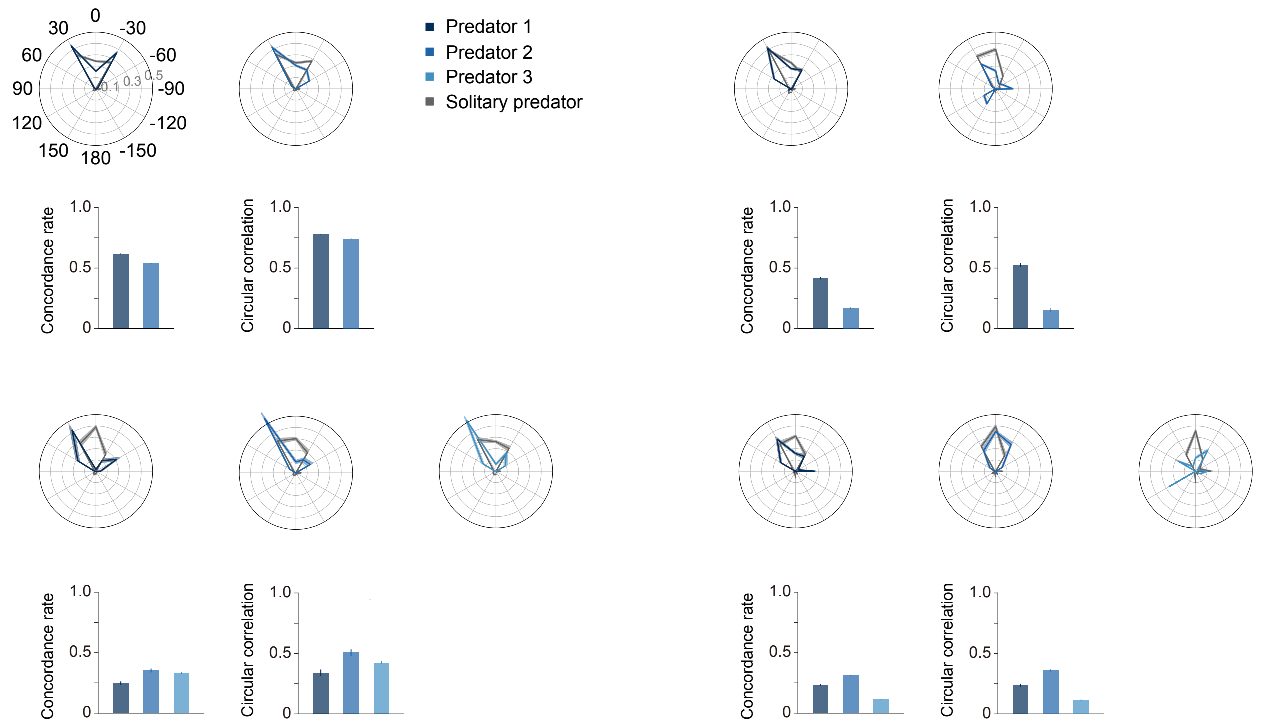


**Figure S2.** Circular histogram, concordance rate, and circular correlation. To visualize the association between each predator in the two- and three-predator conditions and the baseline, which is the predator in the one-predator condition, we made overlays of the frequency of selection for each action. We calculated concordance rates and circular correlations to quantitatively evaluate these associations of action selection. The concordance rate has the advantage of being able to compare all 13 actions, whereas the circular correlation has the advantage of being able to consider the proximity among each action, although it can only evaluate 12 actions, excluding "do nothing". As shown in this figure, the two indices showed similar trends. The predators whose heat maps were similar to that of their prey tended to have higher values on these indices. For all panels, quantitative data denote the mean of 100 episodes ± s.e.m across 10 random seeds.

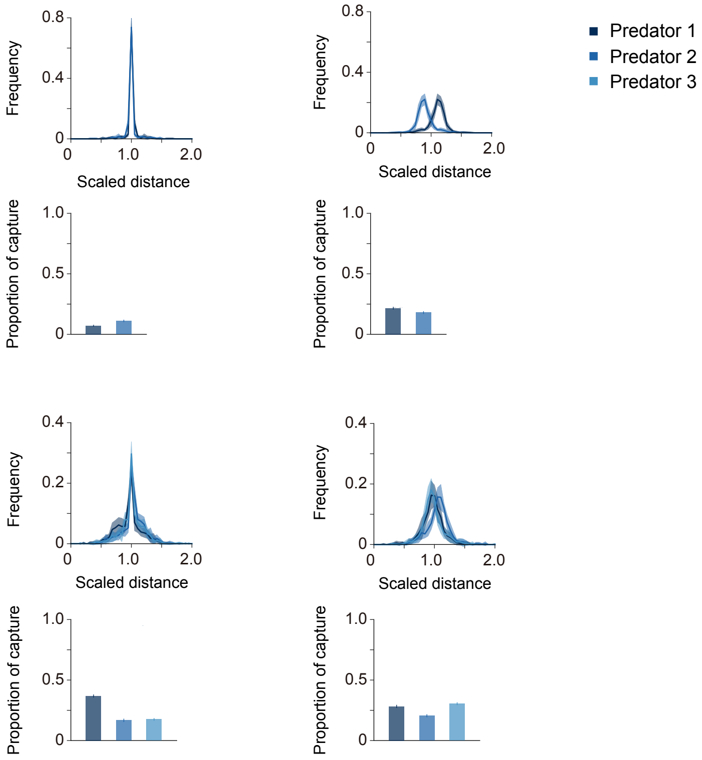


**Figure S3.** Scaled distance among predators and proportion of prey capture in each predator. Scaled distance is a measure of how far a predator moves to capture its prey compared with the other predators during a hunt. Although simplified, this distribution reflects the role of each predator played in a hunt. Specifically, if there is a large difference in the scaled distance among predators, individuals with a larger scaled distance (greater than 1) could play the role of "chaser" (or "driver"), while individuals with a smaller scaled distance (less than 1) could play the role of "blocker" (or "ambusher"). Moreover, if these distances do not differ among predators (concentrated near 1), it is likely that each predator pursued prey in the same manner, suggesting that there was no role division during the hunt. Furthermore, these distributions can be used to capture the flexibility of role division among predators. That is, if the distributions are separate and they do not overlap, there is a division of roles among predators, and these roles are fixed in any hunt. On the other hand, if the distributions are separate but some of them overlap, it indicates that the roles may have switched across hunts. Our results show that the distribution in the individual condition was concentrated around 1, whereas in the shared condition it was divided among individuals. This means that there was rarely role division among predators in the individual condition, while there was role division in the shared condition. These characteristics were more pronounced in the two-predator condition than in the three-predator condition. Perhaps this depends on the episode duration; the duration tends to be longer in the two-predator condition, and the difference in distance is likely to be clearer. Note that even under the two-predator condition, there was some overlap in the distribution in the shared condition. This indicates that the basic roles were fixed among individuals, but interchanged according to the situation (or episode) in the condition. Additionally, because these role divisions are often discussed in the context of cooperation and cheating, we calculated the proportion of prey capture in each predator. The results did not indicate which role was more likely to catch the prey. That is, in the shared $\times$ two condition, the chaser tended to catch more prey, and, on the other hand, in the shared $\times$ three condition, the blocker tended to catch more prey. For all panels, quantitative data denote the mean of 100 episodes $\pm$ s.e.m across 10 random seeds.

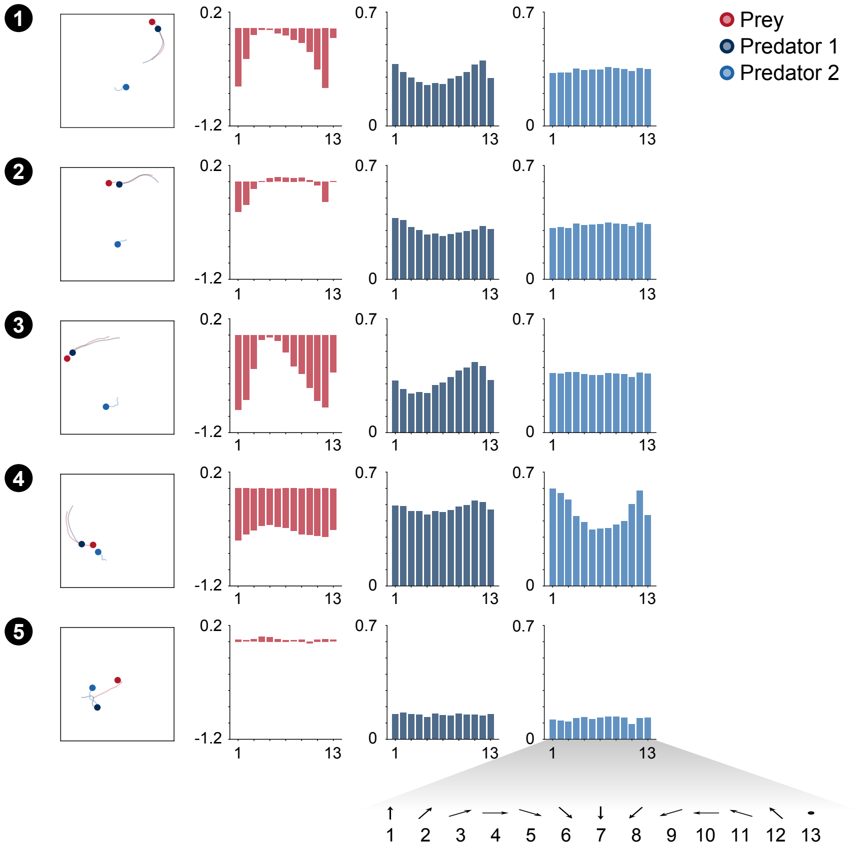


**Figure S4.** Corresponding state-action values (Q-values) for each state. We show the value for each action of each agent in a selected series of state transitions (the scenes 1 to 5). The panels of left side in the figure are the same as the plot in Figure 3. For both predator and prey, each action is defined in terms of a relative coordinate system to the opponent. In other words, action 1 denotes the movement toward the opponent (for the prey, the nearest predator) and action 7 denotes movement in the opposite direction of the opponent. Thus, in the estimated action values of prey, actions 5 to 9 tend to show relatively high values, and in those of predators, actions 1, 2, 3, 11, and 12 tend to show relatively high values.

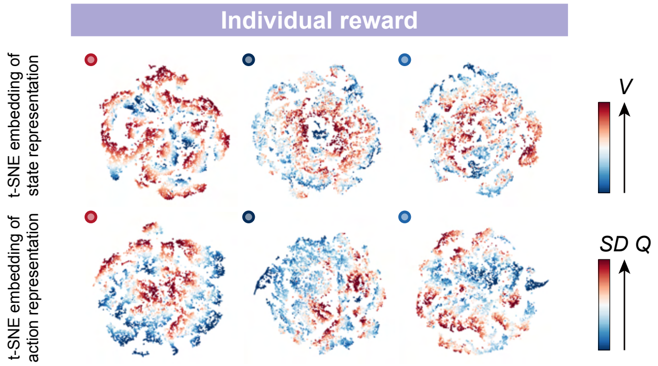


**Figure S5.** Two-dimensional t-SNE embedding of the representations in the last hidden layers of the state-value stream (top) and action-value stream (bottom) in the individual reward condition, in the slow × two condition. The representation is assigned by policy network of each agent to states experienced during predator-prey interactions. The points are colored according to the state values and standard deviation of the action values, respectively, predicted by the policy network (ranging from dark red (high) to dark blue (low)).

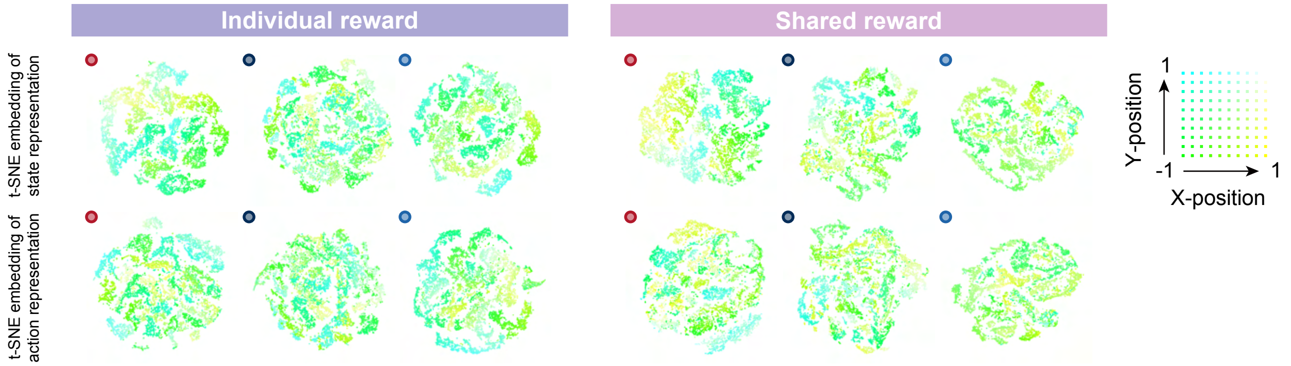


**Figure S6.** Two-dimensional t-SNE embedding colored according to the absolute coordinates of itself in the individual (left) and shared (right) reward conditions, in the slow × two condition. The absolute coordinates (i.e., $x$ and $y$ positions) are directly associated with the reward, as are the distances between prey and predators, because each agent receives a negative reward (-1) for leaving the play area. We therefore colored the internal representation of the agent according to its position. The upper left corner of the play area corresponds to cyan, the lower left to green, the upper right to white, and the lower right to yellow. The embedding of state representations in the prey seems to be roughly clustered according to absolute position, compared to those of the predators. These indicate that prey might estimate the state and action values and make decisions, mediated by absolute position-dependent representations compared to predators.

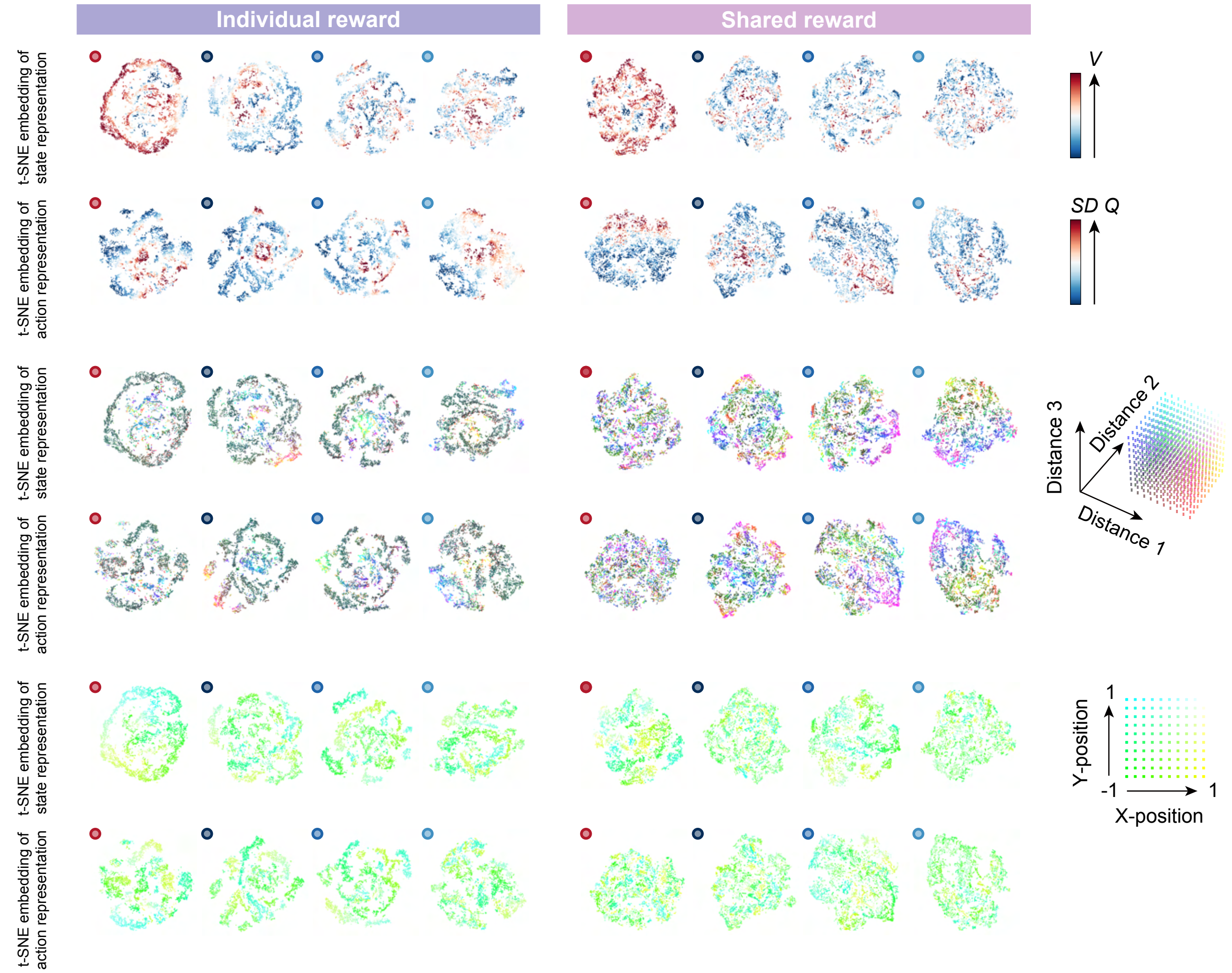


**Figure S7.** Two-dimensional t-SNE embedding of the representations in the last hidden layers of state-value stream and action-value stream, in the slow × three condition. The points are colored according to the state values and standard deviation of the action values predicted by the policy network (top), the distances between prey and predators (middle), and absolute coordinates of itself, respectively.

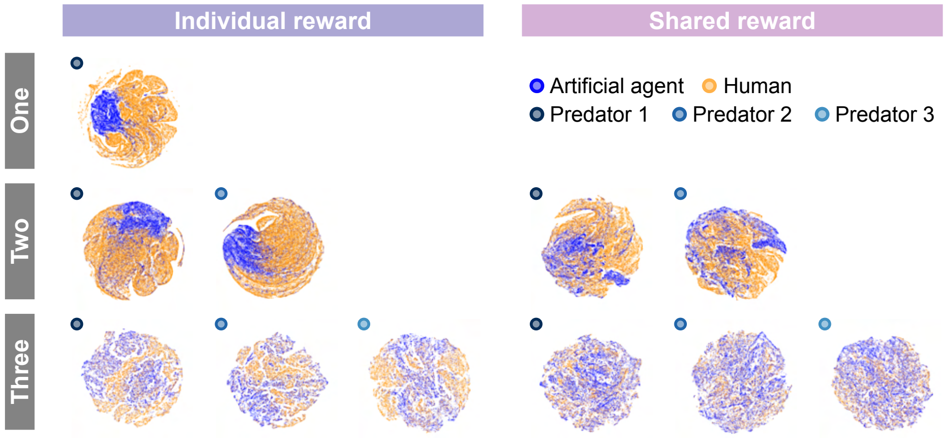


**Figure S8.** Comparison of two-dimensional t-SNE embedding of the internal representations. To visualize the associations of states experienced by predator agents versus agents (self-play) and versus humans (joint play), we show colored two-dimensional t-SNE embedding of the representations in the last hidden layer of the action-value stream. Similar to those of the state stream (Fig. 4b), the experienced states were quite separate, especially in the one- and two-predator conditions.

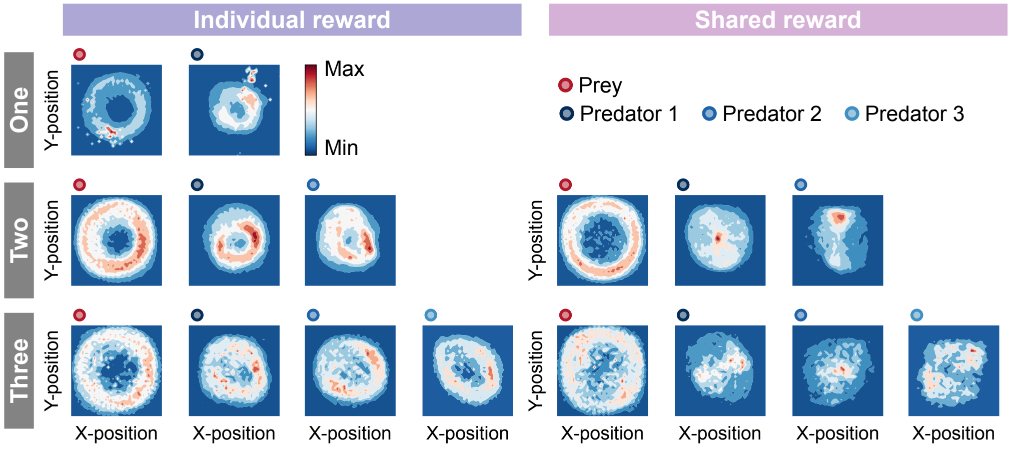


**Figure S9.** Comparison of heat maps between individual (left) and shared (right) reward conditions in joint play. The heat map of each agent was made based on the frequency of stay in each position which is cumulative for 500 episodes (50 episodes $\times$ 10 participants). Our results showed a similar trend between self-play (predator agents vs. prey agent) and joint play (predator agents vs. prey human) in terms of role division among predators. For example, in the individual condition, the heat maps between/among predators were similar, indicating that there was no clear division of roles. On the other hand, in the shared condition, the heat maps between/among predators differed and it indicates that the roles were divided among them. In addition, one of the differences from self-play was the instability of the predator agent's behavior in the one-predator condition. As shown in the figure, under certain conditions, the predator agent stopped moving from its location (the upper right corner of the area).

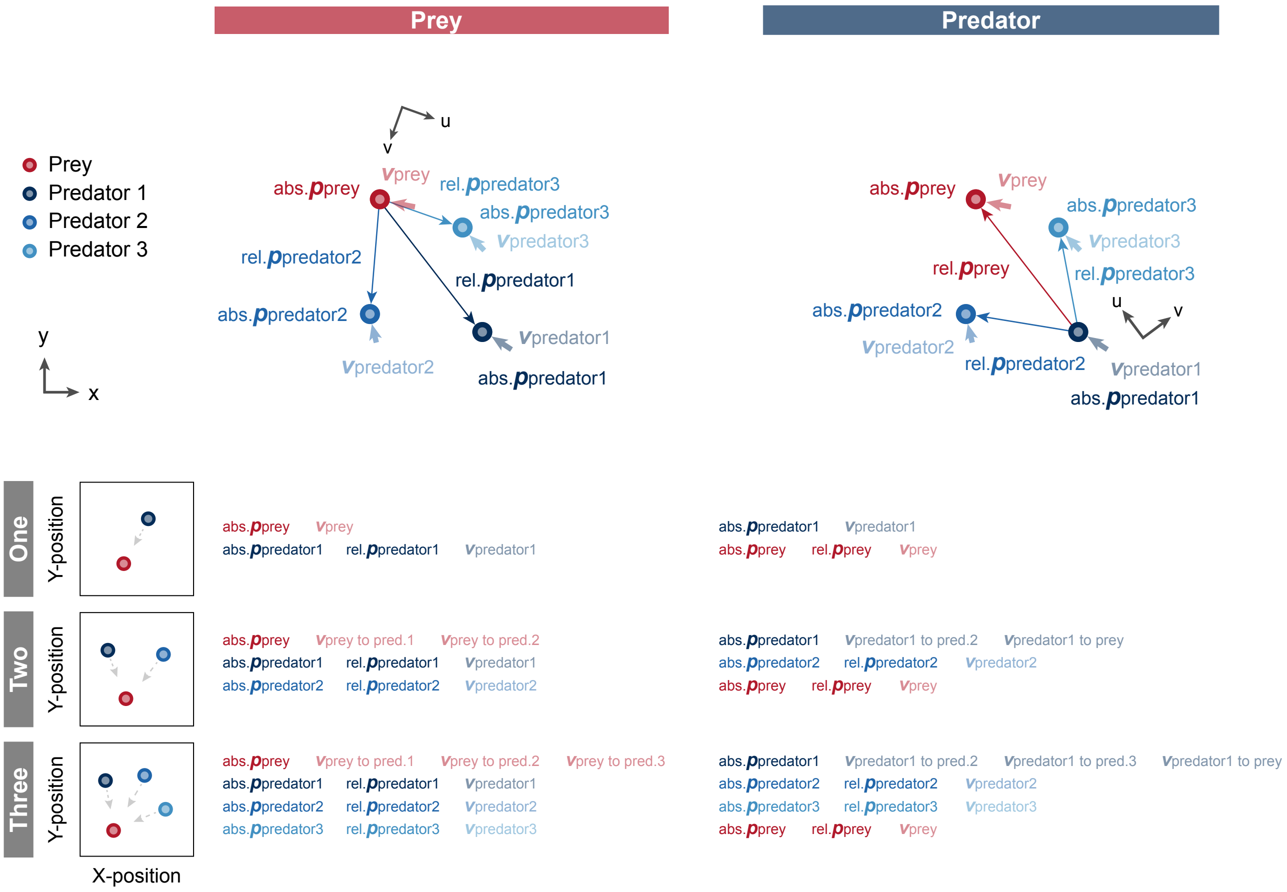


**Figure S10.** Diagram of model input. We used position and velocity information as observation (model input) for each agent. Assuming subjective observation, each variable except absolute position was converted to a relative coordinate system to the opponent, namely, prey for predators and nearest predator for prey, and inputted to the model. Moreover, in the prey input in the three-predator condition, the predator indices were sorted according to the distance between the prey and each predator. Specifically, we considered predator 1, predator 2, and predator 3 in order of decreasing distance. Similarly, in the predator input in the three-predator condition, we set itself as predator 1, the closer predator to itself as predator 2, and the farther one as predator 3. In the figure, abs., rel., bold p, and bold v denote an absolute, relative, position, and velocity, respectively.

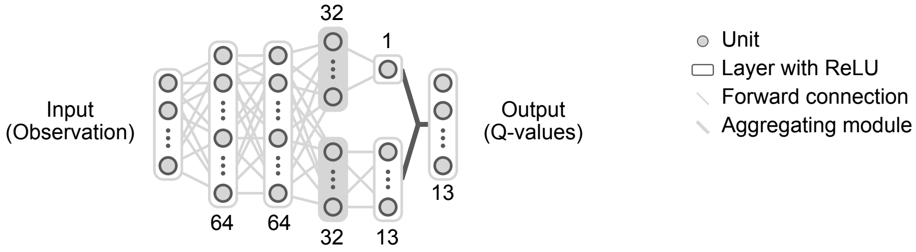


**Figure S11.** Network architecture. The neural network is composed of four layers. The input to the neural network was observation (see Fig. S10) and the output was each value of possible action, namely, a total of 13 action of the 'acceleration' in 12 directions every 30 degrees in the relative coordinate system and 'do nothing'. After the first two hidden layers of the MLP with 64 units, the network branches off into two streams. Each branch has one MLP layer with 32 hidden units. ReLU was used as the activation function for each layer. In the visualization of internal representation of agent, the 32-dimensional hidden vector (parts filled in gray) was embedded in two dimensions, using t-SNE.

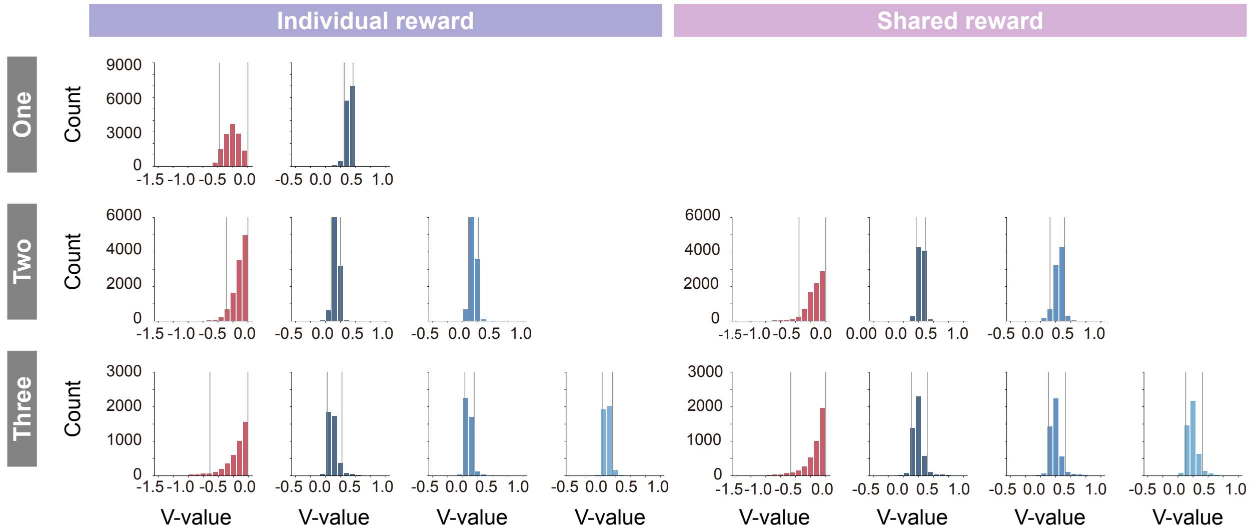


**Figure S12.** Histogram of the state value (V-value) in individual (left) and shared (right) conditions. In coloring the embedding, the lower and upper limits of coloring were set to the 5th percentile and 95th percentile (gray lines), respectively, to prevent visibility from being compromised by extreme values that rarely occur.

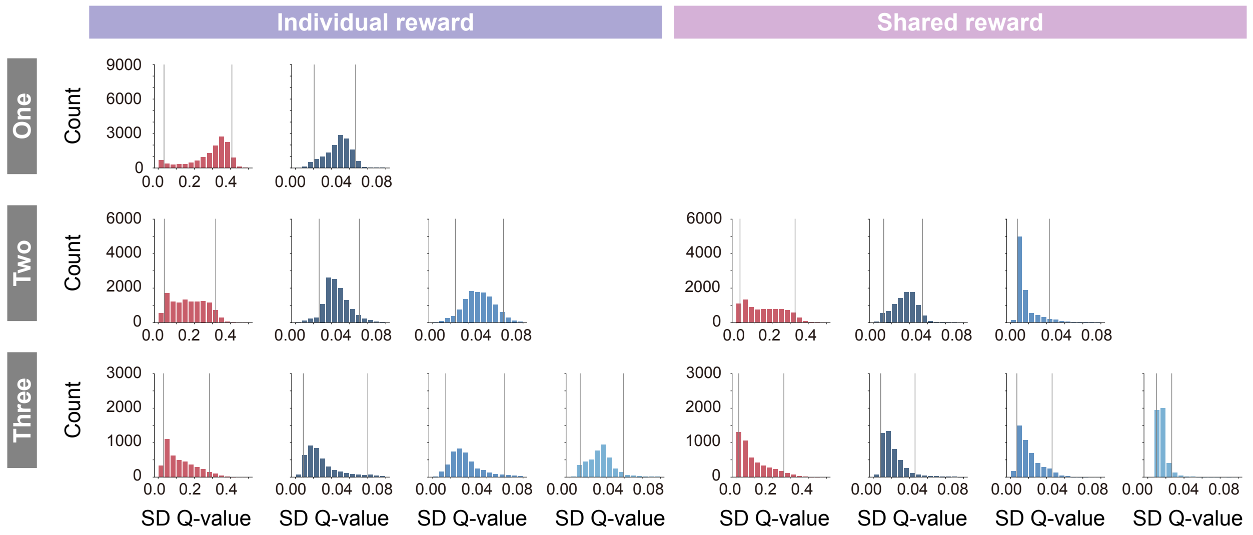


**Figure S13.** Histogram of the standard deviation of state-action value (Q-value) in individual (left) and shared (right) conditions. In coloring the embedding, the lower and upper limits of coloring were set to the 5th percentile and 95th percentile (gray lines), respectively, to prevent visibility from being compromised by extreme values that rarely occur.

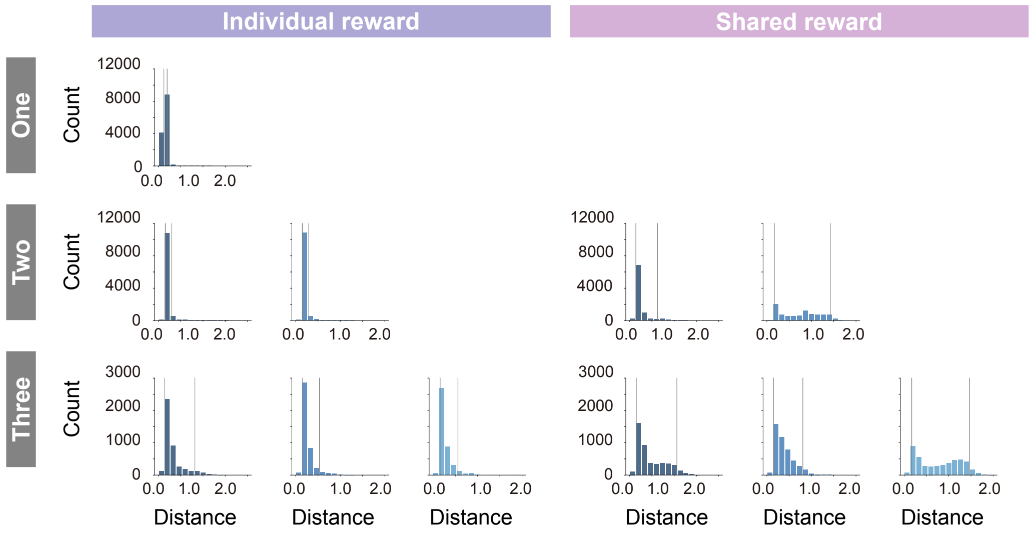


**Figure S14.** Histogram of the distance between prey and each predator in individual (left) and shared (right) conditions. In coloring the embedding, the lower and upper limits of coloring were set to the 5th percentile and 95th percentile (gray lines), respectively, to prevent visibility from being compromised by extreme values that rarely occur.
